# Glial cell activation precedes neurodegeneration in the cerebellar cortex of the YG8-800 murine model of Friedreich’s ataxia

**DOI:** 10.1101/2024.05.17.594658

**Authors:** Andrés Vicente-Acosta, Saúl Herranz-Martín, María Ruth Pazos, Jorge Galán-Cruz, Mario Amores, Frida Loria, Javier Díaz-Nido

## Abstract

Friedreich’s ataxia is a hereditary neurodegenerative disorder resulting from reduced levels of the protein frataxin due to an expanded GAA repeat in the *FXN* gene. This deficiency causes progressive degeneration of specific neuronal populations in the cerebellum and the consequent loss of movement coordination and equilibrium, some of the main symptoms observed in affected individuals. Similar to other neurodegenerative diseases, previous studies suggest that glial cells could be involved in the neurodegenerative process and disease progression in Friedreich’s ataxia.

In this work, we have followed and characterized the progression of changes in the cerebellar cortex of the latest Friedreich’s ataxia humanized mouse model, the YG8-800 (Fxn^null^:YG8s(GAA)>800), which carries a human *FXN* transgene containing more than 800 GAA repeats.

Comparative analyses of behavioral, histopathological, and biochemical parameters were conducted between Y47R control and YG8-800 mice at different time points. Our findings revealed that the YG8-800 mice display an ataxic phenotype, characterized by poor motor coordination, lower body weight, cerebellar atrophy, neuronal loss, and changes in synaptic proteins. Additionally, early activation of glial cells, predominantly astrocytes and microglia, was observed preceding neuronal degeneration along with an increased expression of key pro-inflammatory cytokines and downregulation of neurotrophic factors.

Together, our results show how the YG8-800 mouse model exhibits a stronger phenotype than previous experimental murine models, reliably recapitulating some of the features observed in the human condition. Accordingly, this humanized model could represent a valuable tool to study Friedreich’s ataxia molecular disease mechanisms and for preclinical evaluation of possible therapies.

## Introduction

Friedreich’s ataxia (FRDA, OMIM 229300) is a rare neurodegenerative disease included in the group of neurological disorders called Autosomal Recessive Cerebellar Ataxias (ARCAs). In 1863, the German neurologist Nicolaus Friedreich described the autosomal inheritance pattern of the disease [1], which has a prevalence in Europe of 1-2 cases per 50.000 individuals [2], making it the most frequent of hereditary ataxias. Classically, FRDA has been described as an early onset neurodegenerative pathology, however, symptoms can start at any age, as the clinical evidence has shown the existence of two main drivers of the disease: a developmental hypoplasia that affects the medulla and the spinal cord, and a neurodegenerative process in the cerebellum [3]. The main clinical feature of FRDA is an abnormal development and progressive neurodegeneration of the spinocerebellar system, which in the end is responsible for the progressive loss of movement coordination and equilibrium [4]. Recent findings have described a significant reduction in brain area volumes of the brainstem, superior and inferior cerebellar peduncles, and the dentate nucleus of the cerebellum in people affected by the disease [5]. Moreover, imaging studies have shown white (WM) and gray matter (GM) volume reduction in the cerebellum of FRDA patients [6].

In the majority of cases, FRDA is caused by a mutation in the frataxin (*FXN*) gene consisting of a guanine-adenine-adenine (GAA) expansion that can range between 70 to 1700 repeats or more [7], a number that is directly proportional to symptom onset and severity [8,9]. At the molecular level, this mutation dampens both the transcription rate of the *FXN* gene and the expression of its coded protein frataxin (FXN) through various mechanisms [10,11]. However, the precise molecular mechanism by which GAA triplet expansions reduce *FXN* gene expression remains unclear. FXN is a constitutive protein with several isoforms and subcellular localization [12,13]. The main functions of FXN are related to mitochondrial metabolism and function, and the cellular alterations that occur upon FXN deficiency include deficits in the biosynthesis of iron-sulfur clusters and heme group, iron accumulation and deposition in mitochondria, and increased sensitivity to oxidative stress [14–17]. However, some of FXN functions still need to be clarified, and why the deficiency in this ubiquitous protein affects tissues differently remains unknown.

To date, there is no cure for FRDA. Despite enormous efforts made to develop effective therapeutic strategies, the only treatment for symptom management, Omaveloxolone, was approved by the U.S. Food and Drug Administration in 2023 [18,19]. Therefore, to continue deciphering all the pathological mechanisms underlying the disease and testing other potentially effective drugs, we need to count with *in vitro* and *in vivo* models of the disease that resemble the human pathology as faithfully as possible. Even though several experimental models have been developed, characterized, and successfully used for studying different aspects of FRDA over the last decades, some of them have either a rather mild or a very exaggerated phenotype [20]. Recently, a new model named YG8-800 or YG8JR (Fxn^null^:YG8s(GAA)>800), has been described as a promising humanized model for studying FRDA. The YG8-800 mouse model is based on previous versions of the same model but with a higher number of GAA repeats inserted in the *FXN* gene, which results in even lower FXN levels [20]. Recent reports indicate that affected mice have poor motor coordination reflecting muscular atrophy and signs of cardiac hypertrophy as early as 6 months of age, reduced aconitase activity, and epigenetic changes [21–23]. In this work, we characterized the phenotype of YG8-800 mice and the control strain Y47R (that harbors a normal human *FXN* gene) at the behavioral, histopathological, and biochemical level across different ages. In comparison with control mice, the YG8-800 mice exhibited a clear ataxic phenotype, including a decline in motor performance in the rotarod and foot fault test, less total body weight, cerebellar atrophy, loss of cerebellar granule neurons (CGNs), and synaptic alterations. Notably, we detected an early activation of glial cells, predominantly astrocytes and microglia, preceding neuronal dysfunction, accompanied by increased mRNA transcripts of key pro-inflammatory cytokines in the cerebellum. Our results demonstrate that the YG8-800 mouse model of FRDA reliably mimics the human disease, showing a more severe phenotype than previous FRDA models, with early microglial and astroglial activation, signs of an active neuroinflammatory process.

## Materials and methods

### Animals

All the procedures in this study were approved by the regional animal Ethics Committee of Comunidad de Madrid, Spain (PROEX 013.0/21) and performed in agreement with their guidelines for animal handling and research. Male and female Y47R and YG8-800 mice were purchased from Jackson Laboratory (strains #031007 and #030395, respectively), housed in the animal facility at Centro de Biología Molecular (CBM) Severo Ochoa, with food and water *ad libitum*. Animals were maintained in cages between 20 – 24°C, 55% +/- relative humidity, less than 350 lux and 65 db, and with sensorial enrichments. Only hemizygous Y47R and YG8-800 mice were selected for experimentation, as the latter has lower FXN protein levels and a more severe phenotype than homozygous mice. All animals were weighed every month, between 1 month and 12 months of age. The number of mice/samples used in each set of experiments is specified in each figure and/or its legend.

### Genomic DNA extraction and genotyping

Genomic DNA (gDNA) from hemizygous mice tail biopsies was isolated using the NZY Tissue gDNA isolation kit (NZYTech, Cat. No. MB13503) following the manufacturer’s instructions, using the NanoDrop One/OneC spectrophotometer (ThermoFisher, Cat. No. ND-ONE-W) to determine gDNA concentrations. Mice were genotyped with specific *FXN* primers (see **Additional Table S1**) and conditions described in the quantitative PCR section.

### Behavioural tests

To characterize the ataxic phenotype of animals, we performed the rotarod test and the foot fault test to measure motor coordination, balance, and general locomotion function in mice aged 3, 6, 9, and 12 months.Both sexes were used in a randomized way, without distinction.

#### Rotarod test

The rotarod test was performed as previously described [24,25]. Briefly, mice were placed on the apparatus (Ugo Basile, Italy) for 5 min, recording the time that mice spent on the rotating drum in 2 consecutive trials spaced by 15 min resting periods. We used the accelerating protocol, consisting of acceleration from 4 to 40 rpm during the first 3 min, followed by 2 min at maximum speed. The trial was terminated if mice fell from the apparatus or after a maximum of 5 min, using the best performance (out of two) for quantification.

#### Foot-fault test

For this test, mice were placed on a horizontally elevated (30 cm above the table surface) stainless steel grid of 35 (L) x 40 (W) cm and 1.3 x 1.3 cm mesh size. Animals were left to roam freely on the grid for 1 min, recording their motor activity with a camera placed below the grid, on top of the bench. The scoring system consisted of counting the total number of steps and the number of times the mouse placed a limb through the grid (foot fault). Data are expressed as the percentage of incorrect steps/total steps [25].

### Tissue processing

Animals (1-12 months of age) were deeply anesthetized with isoflurane, followed by transcardiac perfusion with PBS. The brain was immediately removed from the skull, weighed using a precision analytical balance, and measured using a millimetric ruler. Then, it was sectioned into two halves; snap-freezing one half for biochemical studies and post-fixing the other half with 4% PFA for 48h followed by a 10-30% sucrose gradient. To measure the brain size, captured images were quantified using the polygon selection tool of the ImageJ (NIH, Baltimore, USA) software. For immunohistochemical studies, the cerebellum was dissected from hemibrains and embedded in Tissue-Tek O.C.T. (Sakura Finetek Europe, Cat. No. 4583) before being snap-frozen in isopentane (VWR Chemicals, Cat. No. VWRC24872.260). All tissues were stored at -80°C until used.

### RNA extraction and reverse transcription PCR

Total RNA from 3 and 12-month-old mice cerebellum were isolated with the NZY Total RNA Isolation kit (NZYTech, Cat. No. MB13402). To determine RNA purity and concentration we used the NanoDrop One/OneC spectrophotometer (ThermoFisher, Cat. No. ND-ONE-W). Then, the NZY First-Strand cDNA Synthesis Kit (NZYTech, Cat. No. MB12502) was used to perform the reverse transcription polymerase chain reaction (PCR) to obtain complementary DNA (cDNA) sequences, following the manufacturer’s instructions.

### Quantitative PCR

The Luminaris Color HiGreen High ROX qPCR Master Mix kit (ThermoFisher, Cat. No. K0363) was used to perform the quantitative PCR (qPCR) from 12–20ng of cDNA or 10ng of gDNA. The qPCR was performed using the CFX Opus 384 Real-Time PCR System (Biorad, Cat. No. 12011452) and the following program: 10’ at 95°C + 40 cycles of 15’’ at 95°C and 1’ at 60°C + 5’’ at 65°C and increasing 0.5°C until 95°C is reached + 5’’ at 95°C. Intron-spanning primers were used to avoid genomic amplification. The cycle threshold (Ct) of the housekeeping gene *Rn18s* (ribosomal RNA 18S) was used to normalize the Ct of each gene, calculating the relative expression by the 2^-ΔΔCt^ method [26]. Results are expressed as a fold ratio using the data of Y47R mice as control. An additional file specifies all primers used (see **Additional Table S1**).

### Protein extraction and western blotting

Frozen cerebellums from 1, 3, 6, 9 and 12-month-old mice were treated with ice-cold RIPA lysis buffer (50 mM Tris-HCl pH 7.6, 150mM NaCl, 1% Triton X-100, 0.5% sodium deoxycholate, 0.1% sodium dodecyl sulfate (SDS)), 4mM protease inhibitors (Roche Diagnostics, Cat. No. 11697498001) and 1μM okadaic acid (phosphatase inhibitor, Sigma, Cat. No. 459618). Samples were sonicated and centrifuged 10’ at 13,000g at 4°C, and the supernatant containing soluble proteins was stored at -20°C. The BCA kit (ThermoFisher, Cat. No. 23227) was used to determine protein concentration. Samples were denaturalized for 5’ at 100°C using a Laemmli buffer (SDS 10%, β-mercaptoethanol 5%, bromophenol blue 0.5%, and Tris 325mM).

Electrophoresis was performed from 10-20μg of each sample in 4-12% acrylamide-bisacrylamide gradient gels (Invitrogen, Cat. No. NP0322BOX), transferring the samples to nitrocellulose membranes using the iBlot2 dry transfer system (Invitrogen, Cat. No. IB21001). Then, membranes were blocked with 5% non-fat dried milk in PBS and 0.1% Tween-20 (Sigma-Aldrich, Cat. No. 822184) for 60’ and incubated overnight at 4°C with selected antibodies. After washing, they were incubated for 1h at room temperature with peroxidase-conjugated secondary antibodies, using the Amersham ECL prime detection system (GE Healthcare Life Science, Cat. No. RPN2232) to visualize protein bands. ImageJ software was used for signal quantification and values were normalized to the intensity of the loading control GAPDH. All data were normalized to 3-month-old Y47R mice and are expressed as percentages.

Primary antibodies used include rabbit anti-Aconitase 2 (1:1000; Abcam, Cat. No. ab71440), rabbit anti-BDNF (1:1000; Abcam, Cat. No. ab108319), rabbit anti-C3 (1:500; Abcam, Cat. No. ab97462), rabbit anti-Calbindin D28K (1:1000; Chemicon, Cat. No. ab1778), rat anti-CD68 (1:500; BioRad, Cat. No. MCA1957), total OXPHOS rodent WB antibody cocktail (1:500; Abcam, Cat. No. ab23707), mouse anti-FXN (1:1000; Abcam, Cat. No. ab110328), mouse anti-GAPDH (1:5000; Abcam, Cat. No. ab8245), rabbit anti-GFAP (1:1000; Promega, Cat. No. G560A), rabbit anti-GPX4 (1:1000; Abcam, Cat. No. ab125066), rabbit anti-IBA1 (1:1000; Wako Chemicals, Cat. No. 019-19741), rabbit anti-MX1 (1:1000; ThermoFisher, Cat. No. PA5- 22101), mouse anti-NeuN (1:1000; Chemicon, Cat. No. MAB377), mouse anti-PSD95 (1:1000; EMD Millipore, Cat. No. MABN68), rabbit anti-Synaptophysin (1:1000; Synaptic Systems, Cat. No. 1010002), mouse anti-TMEM119 (1:1000; Abcam, Cat. No. ab209064), rabbit anti-TRKB (1:1000; Cell Signalling, Cat. No. 4619P), rabbit anti-pTRKB-Y705 (1:1000; Genscript, Cat. No. A01186).

### Iron content quantification

For iron content measurements, the extracted cerebellum of 3, 6, 9, and 12 months of age animals were sliced with a surgical blade into equal-sized fragments and weighted with a precision balance. Tissue ferrous (Fe^2+^) and total iron content were determined using the colorimetric Iron Assay Kit (Abcam, Cat. No. ab83366) following the manufacturer’s instructions. This kit allows the determination of Fe^2+^ in a sample by generating a coloured complex whose absorbance can be measured at 593 nm. The total iron content can be determined by reducing ferric iron (Fe^3+^) to Fe^2+^ after dissociating Fe^3+^ from its carrier proteins. The absorbance was determined using a Dynex Opsys MR microplate reader (Dynex technologies) and was normalized by mg of tissue. The results are expressed as a percentage of the total iron content of Y47R mice matched for age.

### Histological analyses

The cerebellum of mice aged 3, 6, 9, and 12 months was cryosectioned into 10 μm thick sequential sagittal sections and mounted onto Vectabond^®^-coated (Vector laboratories, Newark, USA) slides. Five consecutive sections corresponding to plate 106 of the Paxinos’s atlas [27] were selected for analysis. Both Nissl and immunohistochemical staining were analyzed by representative photographs acquired with an upright microscope (Nikon 90i, Nikon, Tokyo, Japan) and a DXM1200F camera. The images were acquired and coded for analysis by investigators blinded to the experimental group, analyzing them with the Image J software.

#### Nissl staining

The Nissl method was used to analyze the size of the cerebellum, as well as the thickness of the different layers. Briefly, slides were immersed in 0.25% Toluidine blue (Sigma-Aldrich, Cat. No. 89640) for 30 s, followed by 10 sec rinses in distilled water, dehydrated for 10 sec in 70% EtOH, 10 sec in 96% EtOH, 60 sec in 100% EtOH and cleared in acetic acid/xylene (1:250) for 5 min. Slides were mounted with CV Mount (Leica, Cat. No. 140464300). Low magnification (1x) images were used for analysis, employing the polygon selection tool of ImageJ (Area selection tools). Purkinje cell number was manually counted using photographs taken at 10x magnification, and results were normalized to the length of the corresponding cerebellar folia measured using the ImageJ software.

#### Immunostaining

For semi-quantification of GFAP, as a specific marker of astrocytes; ionized calcium-binding IBA-1, as a marker of microglia; and NeuN, as a marker of cerebellar granule neurons, we used immunostaining methods. Briefly, sections were allowed to warm and then post-fixed in cold 4% PFA for 15 min. After extensive washing in PBS, they were incubated overnight at 4°C with the respective primary antibodies: (i) polyclonal rabbit anti-GFAP (1:2000; Dako, Cat. No. Z0334); (ii) polyclonal rabbit anti-IBA-1 (1:500; Wako Chemicals, Cat. No. 019-19741) and, (iii) monoclonal mouse anti-NeuN (1:5000; Sigma-Aldrich, Cat. No. MAB377). Sections were rewashed with PBS and incubated with the corresponding secondary antibodies: (i) AlexaFluor 488-conjugated secondary anti-rabbit antibody (Invitrogen, Cat. No. A-11008) used at 1:200 for GFAP and IBA-1 (2h at 37°C), rendering green fluorescence and counterstaining the nuclei with DAPI (1:5000; Molecular Probes, Cat. No. 1306) for 10 min; and (ii) EnVision^®^ HRP-labelled polymer, anti-mouse (Dako; Cat. No. K4001, 1 h at RT for NeuN). Finally, the NeuN stained sections were incubated in 3,3′-diaminobenzidine (DAB) substrate-chromogen system (Dako; Cat. No. K3468) to visualize the enzymatic reaction. Representative immunohistochemical photographs for NeuN were acquired at 10x magnification. For fluorescence intensity analysis and quantification, images of 3 leaflets per animal were captured at 20x magnification. The free hand selection tool of the ImageJ software was used to draw regions of interest (ROI) delimiting the total cerebellar area, the molecular layer (ML), the granular layer (GL), and the WM.

### Statistical Analysis

GraphPad Prism 8.00 software was used to perform the statistical analysis. If not specified, data were normalized to the control group Y47R. For statistical comparisons, after outlier identification and removal through the Rout method (Q = 1%), we confirmed if data sets passed the normality D’Agostino-Pearson test. If data sets passed the normality test, two-tailed unpaired Student’s t-tests with a 95% confidence level were used to compare Y47R and YG8-800 mice for each age group. If the normality test was significant, the Mann-Whitneýs U non-parametric test was used. For repeated observations, as in the case of behavioral tests, we performed the mixed effect analysis followed by Sidak’s *post hoc* test. All the statistical details, including the number of animals per group/samples (represented with coloured dots) and the *p* values are specified either in the figure or in the figure legend. Statistical significance was defined when *p* ≤ 0.05.

## RESULTS

### Frataxin-deficient YG8-800 mice show reduced total body weight and progressive motor and coordination impairments

The central nervous system (CNS) is one of the most affected tissues in FRDA. Within the CNS, the cerebellum plays a pivotal role in this disease as evidenced by progressive neurodegeneration and atrophy of the cerebellar cortex GM in FRDA patients [6,28,29]. Therefore, our studies have focused on characterizing cerebellar changes in the YG8-800 mouse model. These mice carry a null mouse *Fxn* allele and the human *FXN* yeast artificial chromosome (YAC) transgene containing around 800 repeats of the GAA trinucleotide expansion. Compared to the Y47R control strain, this experimental mouse model of FRDA has drastically reduced FXN mRNA expression (**Fig. 1A**) and protein levels (**Fig. 1B**) in the cerebellum at all the examined ages. Furthermore, YG8-800 mice exhibit progressive hair loss, reduced body weight, and evident motor impairments (**Fig. 1C, Additional file S2**). Indeed, a significant body weight difference was observed between Y47R and YG8-800 mice, characterized by a reduced body weight in YG8-800 mice since 1 month of age. This difference steadily increases over time evidencing a progressive worsening of the phenotype (**Fig. 1C**).

**Figure 1.**
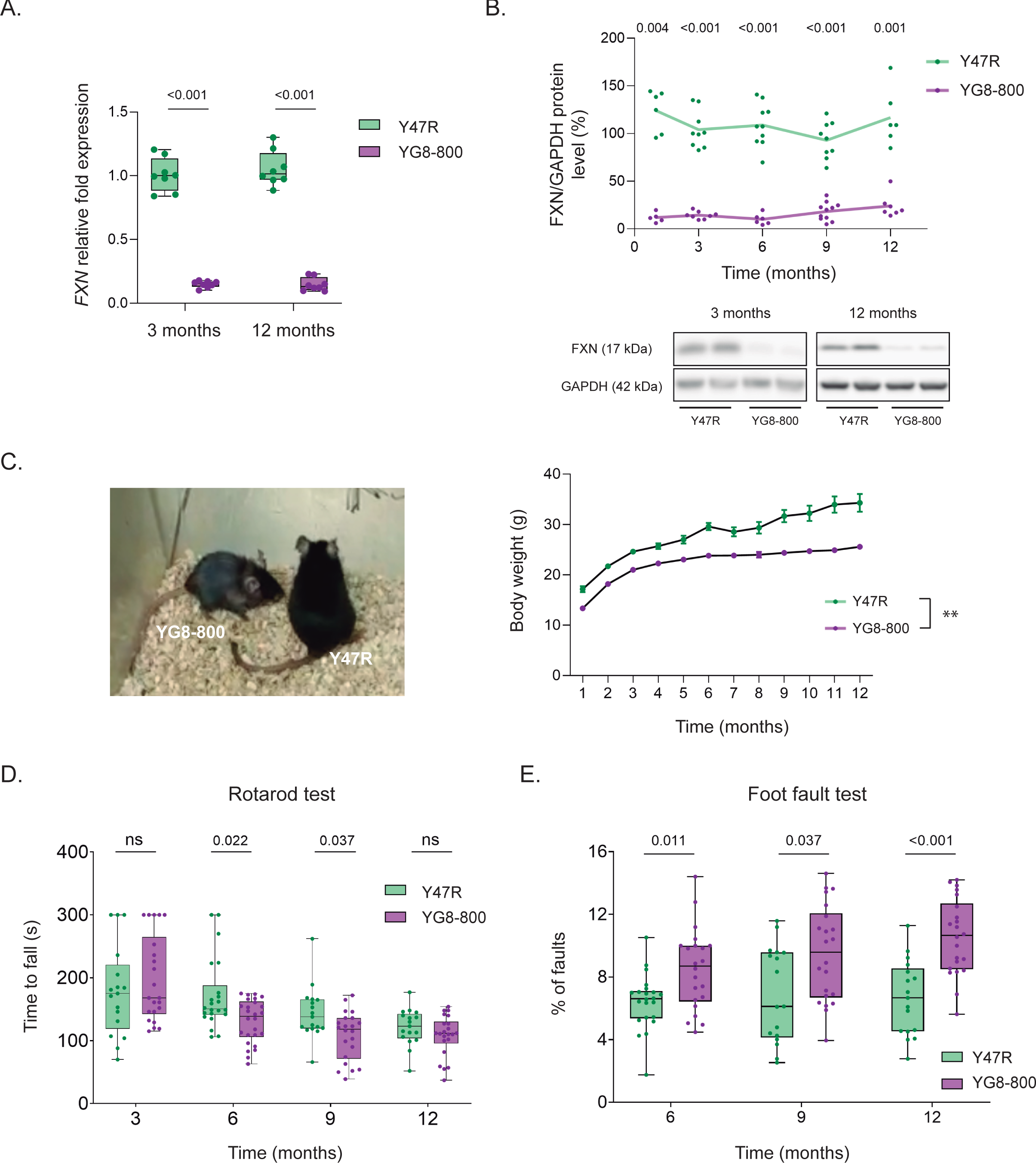
FXN levels and locomotor function in Y47R and YG8-800 mice. A. *FXN* mRNA expression levels of Y47R and YG8-800 mice cerebellum at the indicated months of age determined by qPCR. Gene expression was normalized to the housekeeping gene *Rn18S* and quantified by the comparative Ct method. **B.** Quantification and representative immunoblots of total FXN levels in the cerebellum of Y47R and YG8-800 mice aged 1, 3, 6, 9, and 12 months (data were normalized to the Y47R group, using GAPDH as loading control). The dots represent biological replicates and the connecting lines show the mean value per group at each time point **C.** The image on the left shows the differences in body size between 6-month-old Y47R and YG8-800 animals. The graph on the right compares the total body weight and gain over time of Y47R (n=24-41) and YG8-800 (32-49) mice. **D.** Rotarod test scores in 3-12 months of age Y47R (n=17-22) and YG8-800 (n=22-26) mice. **E.** Foot fault test analysis of 6- 12 months of age Y47R (n=17-22) and YG8-800 mice (n=22). In (**C**) the data represent the mean ± the standard error of the mean.**p<0.01 Vs age-matched Y47R. ns, not significant.

The cerebellum is involved in coordination and motor function; hence, we decided to assess the locomotor function in YG8-800 and age-matched Y47R mice. For this, animals were subjected to a set of behavioral tests, including the accelerating rotarod and the foot fault test. Mouse locomotor performance and coordination were primarily evaluated through the accelerating rotarod assay, starting with mice aged 3 months. When compared to age-matched Y47R mice, the first signs of a statistically significant motor decline were observed when YG8-800 animals were 6-month-old, and those differences were maintained at 9 months of age (**Fig. 1D**). However, at later stages, we did not find significant differences in motor coordination between the two groups. At 12 months of age, the rotarod performance in Y47R mice declined, most probably due to their body weight gain (**Fig. 1D**). To thoroughly evaluate motor coordination, the foot fault test was carried out from 6 months of age onwards. This test quantifies the percentage of paw misplacements/total steps of mice freely moving on an elevated grid. Compared to age-matched Y47R, YG8-800 mice committed significantly more mistakes at 6, 9, and 12 months of age, evidencing that motor coordination was affected in FXN-deficient mice (**Fig. 1E**). Altogether, our results show that the YG8-800 mouse model presents progressive motor impairment and coordination deficits in animals older than 6 months of age.

### YG8-800 mice have cerebellar atrophy and a progressive loss of neuronal and synaptic markers

To detect and follow the progression of gross morphological changes in YG8-800 and Y47R control mice, we measured the total area and weight of the brain every three months between 3-12 months of age. Our observations revealed a significant reduction in the brain weight and area of FXN-deficient mice, which remained consistent over time (**Fig. 2A**). When we assessed the cerebellum area, we observed that, although there was a considerable reduction in the size of YG8-800 mice cerebella compared to the control mice, this reduction was more pronounced as YG8-800 animals aged (**Fig. 2A**). The cerebellum/brain area ratio also showed a progressive reduction in the size of the cerebellum of YG8-800 mice over time.

**Figure 2.**
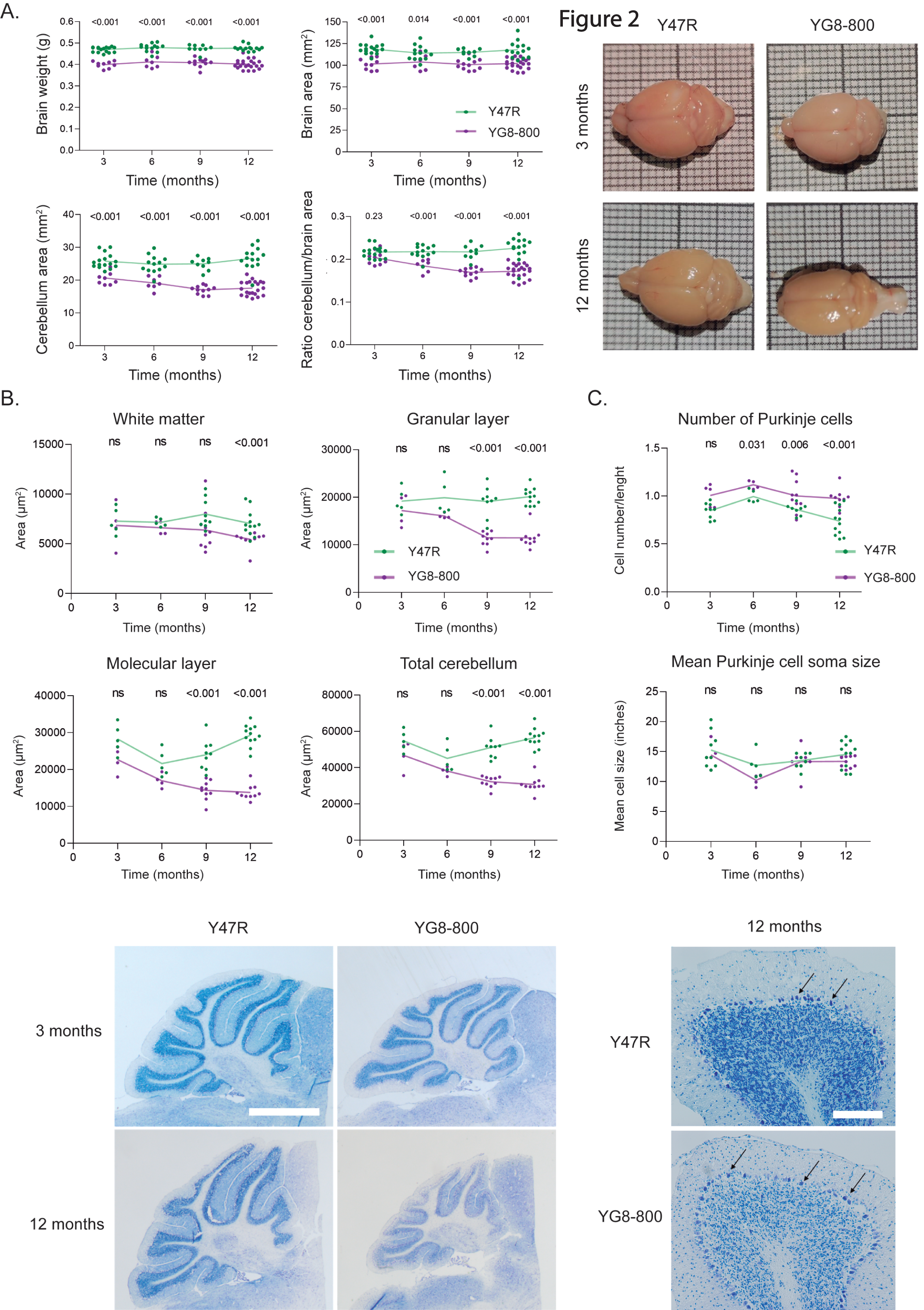
Brain and cerebellum size of the Y47R and YG8-800 mice. A. On the left, graphs show the quantification of the brain weight and area, the cerebellum area, and the cerebellum/brain area ratio of Y47R and YG8-800 mice at different time points. On the right, representative images of Y47R and YG8-800 mouse brains at the indicated time points. The small squares are 1 mm. **B**. Total area of the different cerebellar layers quantified from Nissl staining images of Y47R and YG8-800 mice. The bottom panels show representative white- field images of 3 and 12-month-old Y47R and YG8-800 Nissl stained cerebellums. Scale bar: 1 mm. **C.** The top graphs show the quantification of the number and size of PCs calculated from the Nissl staining images of Y47R and YG8-800 mice cerebellums. On the bottom, representative white-field images of Y47R and YG8-800 Nissl-stained cerebellums. The arrows point to PC somas. Scale bar: 200 μm. In (**A-C**) the dots represent biological replicates and the connecting lines show the mean value per group at each time point. ns, not significant.

When we analyzed the total area of the cerebellum, its different layers (GL and ML), and the WM in Nissl-stained sagittal slices of the two mice strains at different ages, we could determine that the progressive mass loss starts between the ages of 6-9 months in YG8-800 mice. This is mainly due to a reduction in the thickness of the GL and the ML, where the soma of the CGNs and the dendrites of Purkinje cells (PCs) are located respectively (**Fig. 2B**). Interestingly, although the ML of FXN-deficient mice is thinner than in Y47R mice, when the number of PCs was quantified, we detected that YG8-800 animals have more PCs than age-matched control mice, with no differences in the mean cell size between the groups at any of the analyzed ages (**Fig. 2C**). These results suggest that the reduction in the thickness of the ML is probably due to a progressive loss or degeneration of PC dendrites and/or CGN axons, rather than a death of PCs.

To further assess the reduction in the size of the cerebellar cortex of FXN-deficient YG8-800 mice, we evaluated the protein levels of NeuN and calbindin D28K, which are markers for the main excitatory and inhibitory neurons in this brain region; the CGNs and the PCs, respectively [30,31]. Immunoblot results showed a significant and gradual decrease in NeuN levels in these mice at all examined ages, and a progressive slight increase of calbindin D28K levels starting at 9 months of age (**Fig. 3A**). Furthermore, we also detected a progressive decline in the levels of the pre and post-synaptic markers synaptophysin and PSD-95, which further supports the hypothesis of a progressive neuronal loss in the cerebellum of these mice (**Fig. 3B**). In the case of synaptophysin, lower levels of this protein were detected in YG8-800 mice, with values being statistically significant in all cases except for 6 months. For PDS-95, compared to age-matched controls, significantly lower levels were observed in YG8-800 animals that were 6 months old and older. Immunohistochemical analysis of NeuN staining in the cerebellar cortex confirmed a considerable decline in CGN content in 12-month-old YG8-800 mice, compared with Y47R mice (**Fig. 3C**). This reduction in NeuN correlates with the thinning of the GL previously observed with the Nissl staining (**Fig. 2B**).

**Figure 3.**
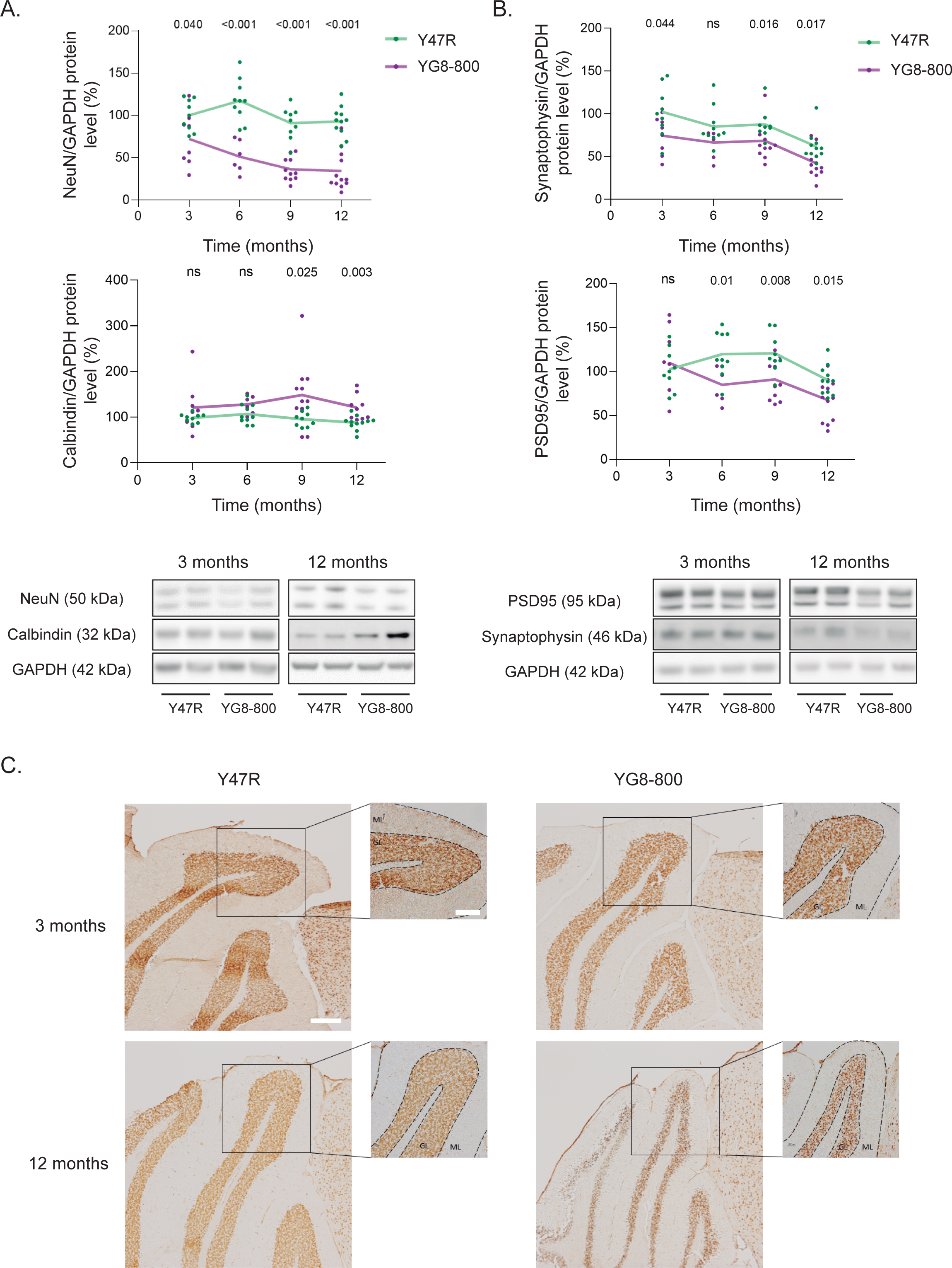
Neuronal markers of CGNs and PCs in the cerebellum of Y47R and YG8-800 mice. A. Quantification and representative immunoblots of NeuN and calbindin D28K protein levels of the indicated strains and months. **B.** Quantification and representative immunoblots of synaptophysin and PSD95 protein levels of the indicated strains and months of age. **C.** Representative images of 3 and 12 months of age Y47R and YG8-800 mice cerebellum sagittal sections stained with the CGN marker NeuN. Scale bars: 250 μm (expanded fields) and 200 μm (insets). GL: Granular layer, ML: Molecular layer. In all plots, data were normalized to the Y47R group. In (**A-B**) GAPDH was used as a loading control, the dots represent biological replicates and the connecting lines show the mean value per group at each time point. ns, not significant.

### FXN deficiency alters iron-sulfur cluster proteins and iron content in the cerebellum of YG8-800 mice

FXN is a protein involved in a wide range of biological processes, participating in iron storage and the formation, repair, and transfer of iron-sulfur clusters (ISC) [32]. These ISC complexes are essential for the correct functioning of numerous enzymes involved in ATP generation, the Krebs cycle, or DNA synthesis, maintenance, and repair [33]. Therefore, we evaluated protein levels of several ISC-containing proteins, including aconitase 2 and CI and CII of the electron transport chain (ETC), in cerebellar samples of YG8-800 and Y47R mice aged 3, 6, 9, and 12 months. At 3 months of age, we did not find significant differences between the genotypes in any of the three proteins evaluated. However, compared to Y47R mice, CI and CII levels were significantly diminished in 6, 9, and 12-month-old YG8-800 mice and aconitase 2 was drastically reduced only in animals aged 12 months (**Fig. 4A**). Additionally, given that iron accumulation has been described in the cerebellum of FRDA patients [34], we measured iron levels in the cerebellum of control and age-matched FXN-deficient mice. Our findings indicate that at 3 months of age, iron levels were similar between YG8-800 and Y47R mice (**Fig. 4B**). By the time that mice reached 6 months of age iron was substantially accumulated in the cerebellum of YG8-800 mice, predominantly the Fe^3+^ form (**Fig. 4B**). The difference in iron content between strains continued as animals aged. To investigate whether this iron accumulation could be linked to ferroptosis, a specific iron-dependent cell death mechanism [35], we assessed the levels of glutathione peroxidase 4 (GPX4), whose activity is normally reduced when ferroptosis is elevated [36]. Our analysis revealed that GPX4 levels remained unchanged between groups at all the ages measured (**Fig. 4C**). Collectively, these results suggest that YG8-800 mice exhibit impairments in the expression of mitochondrial proteins, evidenced by reduced levels of ISC-containing proteins, accompanied by a significant iron accumulation without signs of increased ferroptosis in the cerebellum.

**Figure 4.**
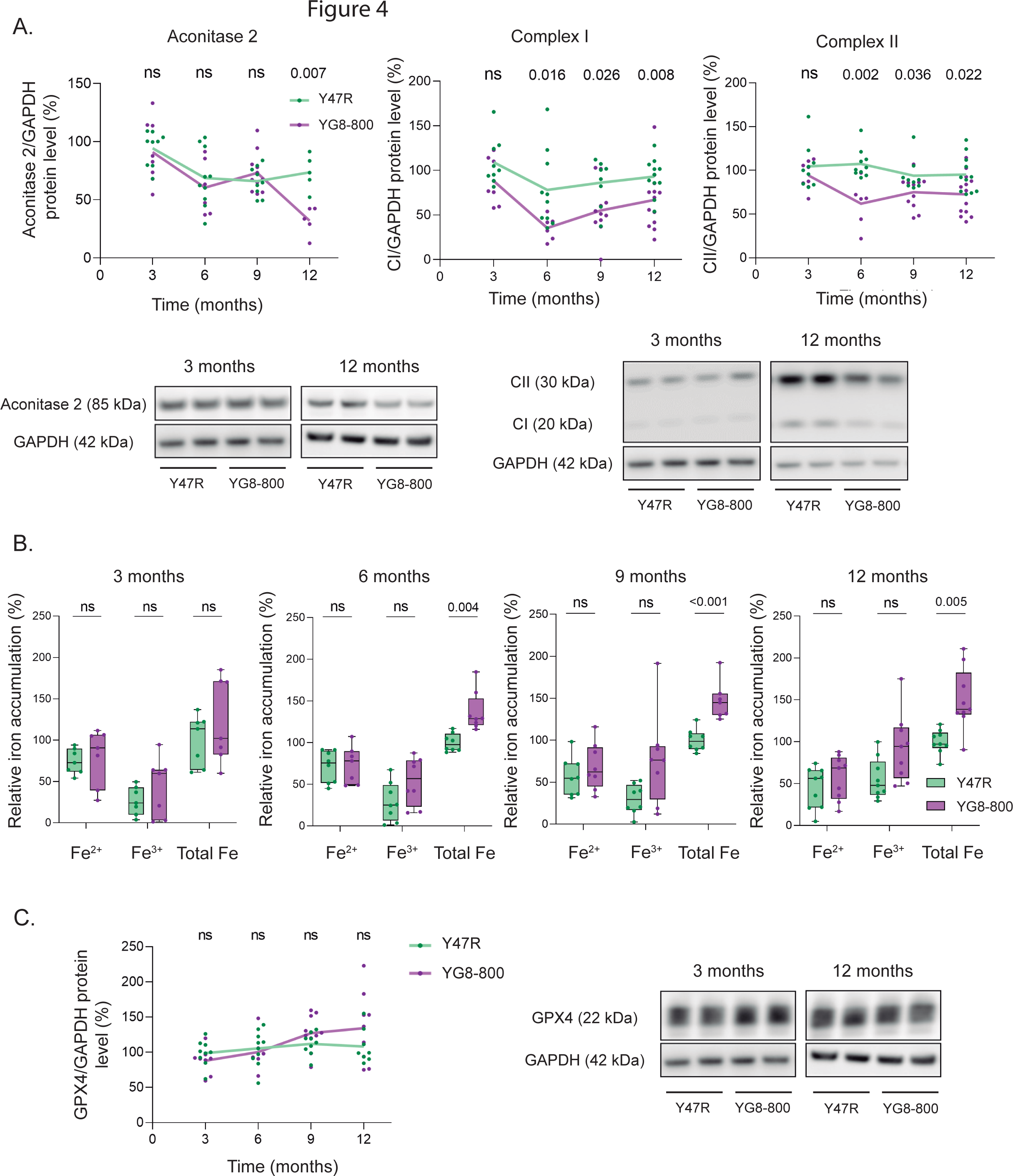
Analysis of iron-sulfur cluster proteins and iron content in the cerebellum of Y47R and YG8-800 mice. A. Quantification and representative immunoblots showing aconitase 2, CI, and CII protein levels in the cerebellum of Y47R and YG8-800 mice at the indicated months of age. **B.** Analysis of Fe^2+^, Fe^3+^ and total Fe content in the cerebellum of 3, 6, 9, and 12 months of age Y47R and YG8-800 mice. **C.** Graph and representative immunoblots showing the quantification of GPX4 protein levels at the indicated months of age. In (**A, C**) GAPDH was used as a loading control, normalizing the data to the Y47R group. The dots represent biological replicates and the connecting lines show the mean value per group at each time point. ns, not significant.

### Glial cell reactivity and neuroinflammation precede neurodegeneration in the YG8- 800 mice cerebellum

Glial activation/reactivity are common features observed in different animal models and tissues from patients suffering from multiple neurodegenerative diseases. Therefore, we have characterized the activation/reactivity status of microglia and astrocytes in the cerebellum of YG8-800 and Y47R control mice over time (from 1 to 12 months of age). Our analysis of different markers of activated microglia revealed that IBA1 levels in YG8-8000 mice were significantly higher than those of age-matched Y47R animals since they had 3 months of age (**Fig. 5A**). Regarding the proinflammatory markers CD68 and TMEM119, there is also an increase in the levels of these two proteins in the cerebellum of FXN-deficient mice. Still, values only reached statistical significance at later stages of the disease, i.e. at 12 months of age for CD68, and 9 months for TMEM119 (**Fig. 5A**). In addition, we assessed IBA1 immunoreactivity in the cerebellar cortex and WM. Our analysis showed consistently elevated fluorescence intensity levels of IBA1 at every examined age in YG8-800 mice relative to control mice (**Fig. 5B**). Notably, significant differences in IBA1 immunofluorescence intensity were mostly restricted to the GL of YG8-800 animals, by 9 months of age the increased levels of IBA1 extended to the ML, reaching the WM when animals were 12-month-old.

**Figure 5.**
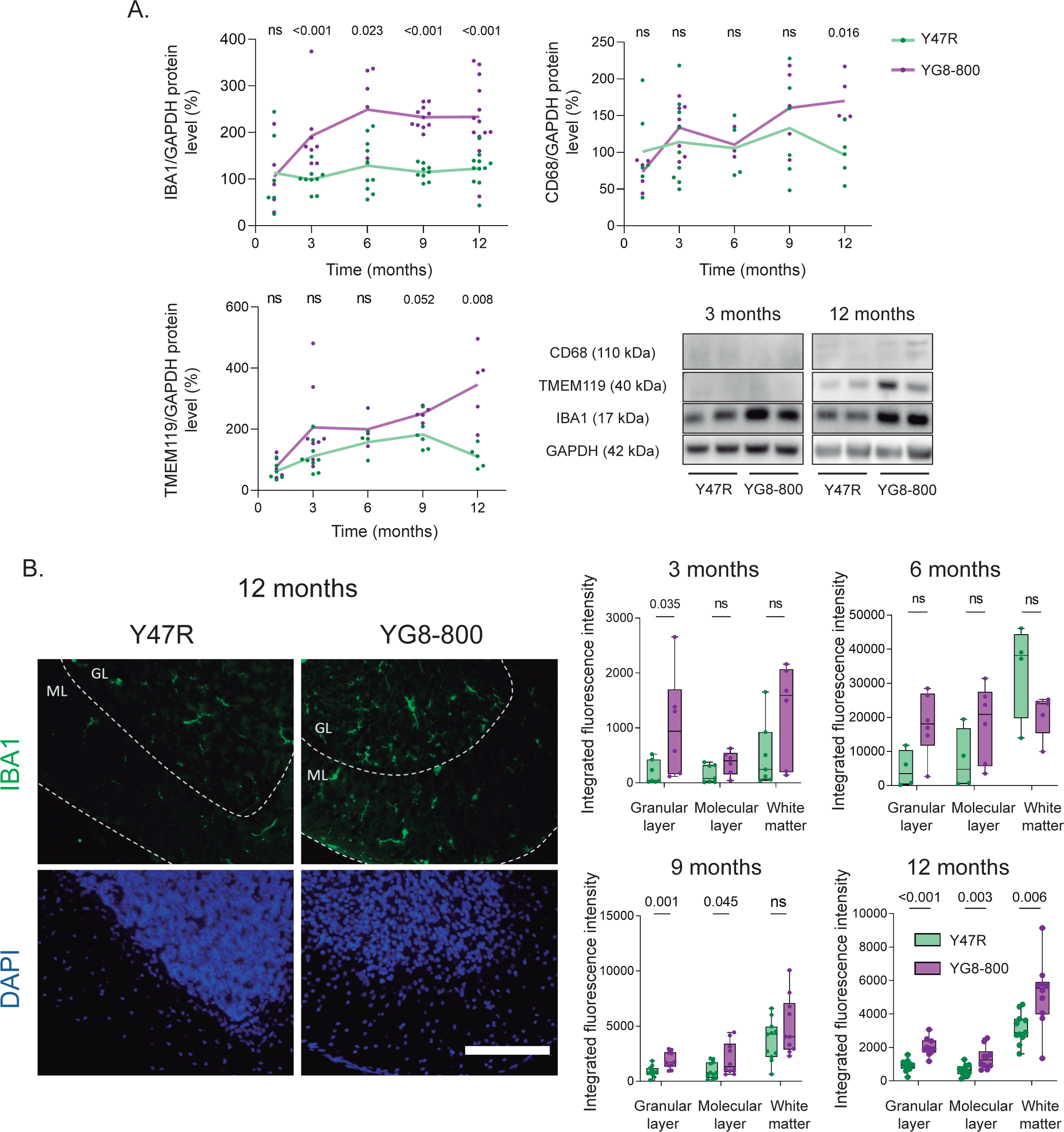
Microglial activation in the cerebellum of Y47R and YG8-800 mice. A. Quantification and representative immunoblots of IBA1, CD68, and TMEM119 protein levels in both mice strains at the indicated months of age (data were normalized to the Y47R group, using GAPDH as loading control). The dots represent biological replicates and the connecting lines show the mean value per group at each time point. **B.** On the left, representative photomicrographs of 12 months of age Y47R and YG8-800 mice cerebellar slices labeled with the microglial marker IBA1 (green) and DAPI to stain the nuclei (blue). Scale bar: 200 μm. GL: Granular layer, ML: Molecular layer. On the right, quantification of IBA1 fluorescence intensity in the different cerebellar layers and the white matter of Y47R and YG8-800 mice at the indicated months of age. ns, not significant.

Regarding astrocyte reactivity, similar to what we found in microglia, immunoblot results showed a progressive increase in protein levels of the main activation marker GFAP in the cerebellum of FXN-deficient mice. This increase was already significant in the cerebellum of 1-month-old YG8-800 mice. Values remained higher than those of control mice even at 12 months of age (**Fig. 6A**). Along with higher levels of GFAP, we detected a progressive increase in C3 levels, which is one of the main proteins associated with a neuroinflammatory reactivity phenotype (**Fig. 6A**) [37]. A similar trend was observed for MX1, another marker of astroglial reactivity. The levels of this protein progressively increased as YG8-800 animals aged, reaching a statistical significance as early as 6 months of age (**Fig. 6A**). As in the case of IBA1, when we examined GFAP by immunofluorescence, fluorescence intensity levels in cerebellar sagittal slices were notably higher in YG8-800 than in control mice since 3 months of age (**Fig. 6B**). When intensity levels were separately quantified in the ML, the GL and the WM, GFAP labeling showed a similar pattern to IBA1. Higher GFAP levels were observed in YG8-800 mice first in the GL (starting at 1 month of age) and then extended to the ML (from 9 months onwards). We did not find differences in GFAP values between genotypes in the WM (**Fig. 6B**).

**Figure 6.**
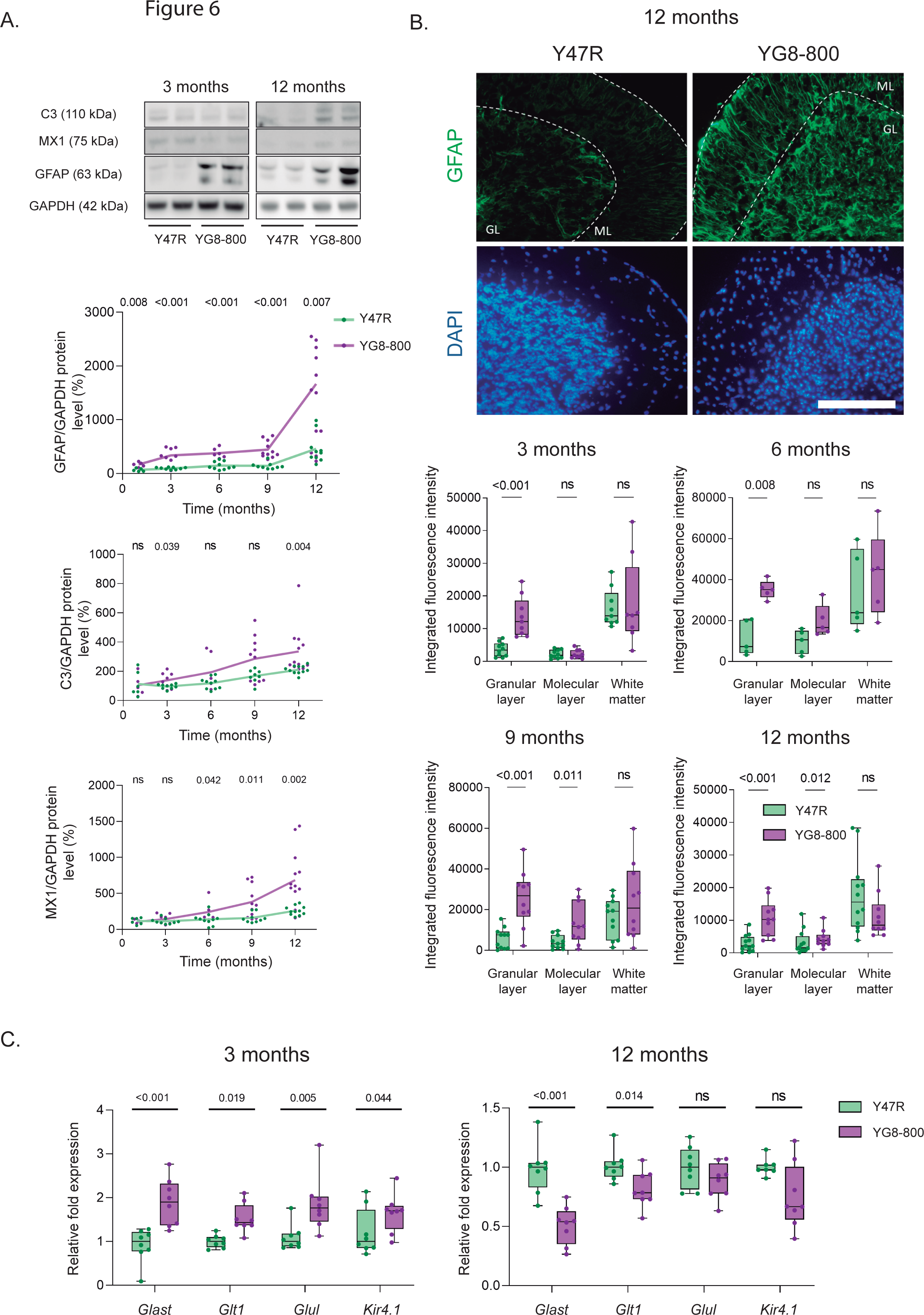
**Astrocyte activation and reactivity markers in the cerebellum of the Y47R and YG8-800 mice**. **A.** Representative immunoblots and quantification of GFAP, C3, and MX1 protein levels at the indicated months of age in both mice genotypes (GAPDH as loading control). The dots represent biological replicates and the connecting lines show the mean value per group at each time point. **B.** On top, representative photomicrographs of Y47R and YG8-800 mice cerebellum at 12 months of age immunostained with specific antibodies against GFAP (green) and DAPI, to stain the nuclei (blue). Scale bar: 200 μm. GL: Granular layer, ML: Molecular layer. The bottom graphs show GFAP fluorescence intensity quantification in the different cerebellar layers of Y47R and YG8JR mice at the indicated months of age. **C.** *Glast*, *Glt-1*, *Glul* and *Kir4.1* mRNA expression levels in the cerebellum of 3 and 12-month-old Y47R and YG8JR mice determined by qPCR. In all cases, gene expression was normalized to the housekeeping gene *Rn18S* and quantified by the comparative Ct method. In all plots, data were normalized to the Y47R group. ns, not significant.

In addition, we evaluated the transcript expression of several proteins involved in astrocytic homeostatic functions, such as the glutamate transporters *Glast* and *Glt-1*, the glutamate synthase enzyme (*Glul),* and the potassium channel *Kir4.1.*, in the cerebellum of both mice strains. The levels of all these transcripts were upregulated in 3-month-old YG8-800 mice, compared to age-matched Y47R mice. By the time the mice reached 12 months of age, the levels of these transcripts either remained unchanged (*Glul* and *Kir4.1*) or were downregulated (*Glast* and *Glt-1*) (**Fig. 6C**).

### The pro-inflammatory/neuroprotective balance in the cerebellum of YG8-800 mice is altered

After observing a strong glial cell activation in the cerebellum of YG8-800 mice at early and late stages of life and disease progression, we analyzed the expression of various genes, including cytokines and neurotrophic factors, involved in the inflammatory response and the neuroprotective response in cerebellum samples of YG8-800 mice and the respective controls. The profile of differentially expressed genes in mice aged 3 and 12 months is shown in heatmaps (**Fig. 7A**). Compared to age-matched control animals, we observed a significant increase in the *iNos/Arg1* ratio in the cerebellum of YG8-800 mice as early as 3 months of age. Moreover, some cytokine transcripts associated with the inflammatory response (*Il1b, Tnfa, Igfbp2*) were upregulated as well in these samples. At this same age, expression of neurotrophins like *Bdnf, Tgfb1,* and *Fgf2* was increased, while Ngf expression decreased, and the rest of the transcripts remained unchanged (**Fig. 7A**).

**Figure 7.**
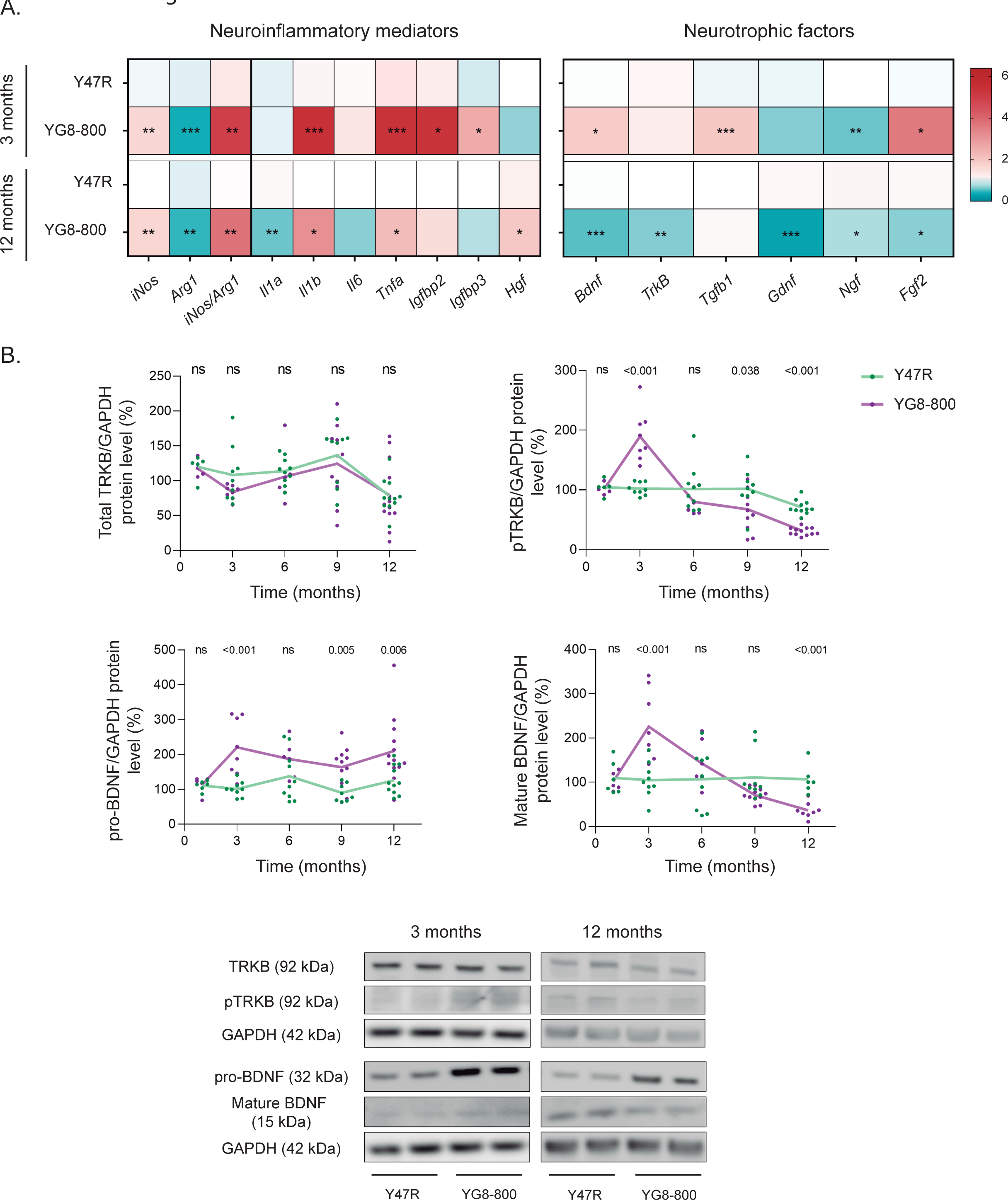
**Analysis of the proinflammatory/neurotrophic balance status in the Y47R and YG8-800 mice cerebellum**. **A.** Heatmap showing mRNA expression levels of pro-inflammatory factors and genes involved in different neurotrophic pathways in the cerebellum of 3 and 12 months of age Y47R and YG8-800 mice determined by qPCR. In all cases, gene expression was normalized to the housekeeping gene *Rn18S* and quantified by the comparative Ct method. *p<0.05, **p<0.01, ***p<0.005 Vs age-matched Y47R. **B.** Quantification and representative immunoblots of total TRKB, pTRKB, pro-BDNF and mature BDNF at the indicated months of age (GAPDH as loading control). The dots represent biological replicates and the connecting lines show the mean value per group at each time point. In all graphs, data were normalized to the Y47R group. ns, not significant.

When we compared the expression profile of the same genes in cerebellum samples of 12- month-old mice of both genotypes, we observed that most of the proinflammatory transcripts that were upregulated at 3 months of age, remained upregulated in samples of YG8-800 mice. On the contrary, at this time point, the expression of the majority of the neurotrophins we examined was significantly downregulated (**Fig. 7A**). These results suggest that a neuroinflammatory component is present from the early stages of the disease, while a deficiency in neurotrophic factors is a progressive process that gradually increases as the pathology advances.

Interestingly, when we evaluated the BDNF signaling pathway in more detail in the cerebellum of 1-12-months-old YG8-800 and Y47R mice, we found that both the mature form of this protein and the activated form of its receptor (pTRKB-Y705) were significantly elevated in YG8-800 mice at 3 months (**Fig**. **7B**). However, these levels gradually declined over time. Furthermore, starting at 3 months of age, protein levels of the precursor form of BDNF (pro- BDNF) were higher in the cerebellum of YG8-800 mice than in control mice (**Fig. 7B**), overall suggesting a progressively hampered activation of the BDNF neuroprotective pathway in FXN-deficient mice as the pathology advances.

## Discussion

FRDA is a neurodegenerative disease for which there is still no cure. Current therapy is based on symptomatic treatments such as the recently FDA-approved compound, Omaveloxolone [38]. This compound reduces inflammation and oxidative stress in FRDA patients by induction of the nuclear factor erythroid 2-related factor 2 (NRF2). It is forecasted to be available in Europe next year, as it was recently approved by the European Medicines Agency. As FRDA is a multifaceted rare disease, a significant effort is being invested to try to find new drugs and therapeutic approaches that could arrest/limit symptoms or could be used as complementary therapies. However, such approaches are developed from knowledge of basic cellular and molecular mechanisms, and still much research is needed to fully understand the pathological mechanisms of FRDA. One of the reasons why this knowledge has lagged is the lack of animal models that could faithfully mimic the clinical and pathophysiological manifestations observed in FRDA patients. During the last decades, different animal models using several approaches have been developed and successfully characterized. However, although many of these models show similarities with the human disease, their phenotype has been determined as either too mild or too exaggerated. The novel YG8-800 mouse is probably the most promising animal model for the study of the FRDA-disease mechanisms up to date. Despite some controversy regarding heart disease in this model, previous studies have shown that these mice have, among others, a behavioral phenotype of varying severity but comparable to movement coordination problems observed in FRDA patients, reduced aconitase activity, and epigenetic changes [21–23,39]. Therefore, in this work, we aimed at characterizing the disease progression regarding the neurodegenerative/inflammatory processes in the cerebellar cortex of these YG8-800 mice, comparing the results with the corresponding control strain.

In line with other studies, we observed that YG8-800 mice have remarkably less *FXN* mRNA and FXN protein levels in the cerebellum, less body weight, and severe hair loss than control mice [21]. Notably, YG8-800 mice showed a phenotype that resembles an ataxic behavior characterized by uncoordinated movements and gait instability. By analyzing locomotor function with the rotarod test, we have shown that YG8-800 mice exhibit deficits in motor coordination that start at 6 months of age, a time point slightly later compared to the findings reported by Kalef-Ezra et al. [23]. Discrepancies in results between studies could be attributed to variations in the experimental protocols with more restrictive and prolonged ages studied, as well as larger sample sizes used in this work. The differences observed at 6 and 9 months of age in the rotarod test between the two genotypes were no longer significant when animals reached 12 months. In our hands, Y47R mice were significantly heavier than their YG8-800 counterparts, and the difference in body weight between groups progressed as animals aged, being drastically different at later time points. According to some studies, higher body weight is correlated with poor rotarod performance [40]. Therefore, the decreasing rotarod performance in Y47R mice may be attributed to their body weight rather than other neurobehavioral factors. Besides, general locomotor function was assessed using the foot fault test [25,41]. Similar to what we found in the rotarod test, we detected differences between Y47R and YG8-800 mice aged 6 months. However, as the results of this test are not affected by body weight, the differences were maintained until later.

Consistently with reduced body weight, the brain and the cerebellum of YG8-800 mice are significantly smaller than those of control animals. Moreover, assessments of the cerebellum/brain ratio indicated that while the difference in the brain size between YG8-800 and Y47R mice remains constant, the cerebellum of YG8-800 mice progressively shrinks as the animal ages, evidencing a neurodegenerative process. Our results suggest that the smaller size of the cerebellum is mainly due to a reduction of the GL and ML, while the WM is not that affected in the slices examined. The reduction of the GL, which starts when YG8-800 mice are between 6 and 9 months old, is very likely due to a progressive loss CGNs, as we have observed a progressive reduction in NeuN labeling, an exclusive marker of CGNs and whose reduced levels are associated with neurodegeneration. Future research should address whether CGN death is due to ferroptosis induced by the observed iron accumulation, by apoptosis, or by any other mechanism. Nevertheless, these results are consistent with magnetic resonance imaging studies in FRDA patients in which a reduction in GM volume in the cerebellum was observed [5]. In contrast with the results of the aforementioned study, our data do not indicate that severe changes are occurring in the WM of the slices analyzed. However, a thorough evaluation of changes in this cerebellar area is needed to determine the damage extension in this FRDA murine model fully.

Of note, we observed that the area of the ML was smaller in YG8-800 mice which seems to be in contrast to having a similar or slightly higher number of PCs. As we have detected less NeuN labeling in the GL of these mice, we cannot rule out that the reduction in the ML could be in part reflecting degeneration of CGN axons located in the GL. Besides, several studies have shown abnormal PC development and impairment of dendritic arborization in different models of ataxia [42]. Since we observed normal PC somas in YG8-800 mice, we may speculate that rather than dying, PC dendritic trees could be degenerating, also contributing to the ML reduction. Further research will help determine if PC dendrites and/or CGN axons are altered in this model of FRDA.

Neuronal loss in specific brain areas is a common feature of spinocerebellar ataxias like FRDA. However, emerging evidence shows that neurodegeneration could result from an inadequate function of the surrounding glial cells, highlighting the significant role that non-neuronal cells such as microglia and astrocytes could have in the pathogenesis and progression of neurodegenerative diseases like FRDA [43–47]. Glial cells are essential cells in charge of maintaining brain homeostasis. Under pathological conditions, these cells activate and establish a positive inflammatory response that helps restore neuronal function. However, an exacerbated inflammatory response can alter glial cell function, promoting a reactivity state of both microglia and astrocytes that could have negative consequences for the neighboring neurons, worsening or accelerating the progression of a pathological condition [48–50]. In our hands, we have observed an early activation of microglia and astrocytes in the cerebellum of YG8-800 mice, evidenced by a significant increase in the levels of IBA1 and GFAP, respectively. These increments are already present in some cases in 1-month-old mice, indicating that these cells acquire an activated state at the early stages of the disease probably due to changes in the cerebellar microenvironment. Besides, accompanying the activation of both types of glial cells, there is a later increase from around 6 to 9 months of age of other markers that indicate an active pro-inflammatory state, i.e. CD68 and TMEM119 for microglia, and C3 and MX1 for astrocytes. Furthermore, in YG8-800 mice we found spatiotemporal heterogeneous patterns consisting of early upregulation and late downregulation of some transcripts that codify proteins associated with essential homeostatic functions in astrocytes. These patterns could be reflecting an early attempt by astrocytes to maintain their homeostatic functions. Still, similar to what happens in other diseases where neuroinflammation becomes chronic, it is possible that in this FRDA mouse model astrocytes are progressively losing those functions, ultimately harming neuronal function and viability.

When we analyzed in more detail the status of activated glial cells in the layers of the cerebellar cortex, we observed that this process started at a different pace but followed a similar pattern for both cell types. In YG8-800 animals, we observed an early activation of astrocytes and microglia in the GL of 3-month-old mice, followed by a subsequent activation in the ML (around 9 months) that eventually reached the WM (around 12 months in microglia). Whether the activation of glial cells in the GL is the cause or the consequence of CGN degeneration remains to be determined. A degenerative process of CGNs, before massive cell death, may release signals triggering the activation of astrocytes and microglia. Alternatively, glial cell activation may release signals promoting the degeneration of CGNs.

Along with a strong glial activation, we detected significant changes in some neuroinflammatory mediators and neurotrophic factors in YG8-800 mice. In 3-month-old animals, many of the pro-inflammatory mediators and some of the neurotrophic factors analyzed were upregulated. However, when mice reached 12 months of age, what would represent later stages of the disease, neuroinflammation was still present in the cerebellum, but the levels of neurotrophic factors significantly dropped down, with BDNF being one of the most notorious examples of this phenomenon. Similar results regarding BDNF have been observed in a transgenic model of spinocerebellar ataxia type 1, which was specifically associated with PC death [51]. In our study, we did not detect PC death, but we observed a similar age-dependent upregulation/downregulation of BDNF, which may be a compensatory mechanism to support CGN survival (at early stages of the pathology) in an autocrine way as this factor is mainly expressed by excitatory neurons [52]. The early increase of BDNF expression in CGNs could be a molecular mechanism triggered to avoid cell death. However, as CGNs progressively die the levels of this neurotrophic factor decrease. Nevertheless, this possibility was not directly addressed in our experiments. Taken together, our observations suggest that besides having changes that possibly affect their functioning, early glial activation in the YG8-800 FRDA murine model is actively contributing to neuroinflammation and possibly to neuronal death. We consider that glial activation precedes neurodegeneration because apart from lower NeuN levels and a slight decrease in synaptophysin, which occur at 3 months, changes in the rest of the parameters associated with neuronal function were detected at later stages of the disease, whereas glial activation was present before. In any case, a more precise characterization of the activity/reactivity profile of these cells is needed. Moreover, it would be interesting to determine if glial activation and neuronal death also occur in other brain areas.

The results of our characterization are in line with previous reports showing that this experimental model of FRDA is a promising model to study disease mechanisms. At the early and late stages of the disease, we observed a clear ataxic phenotype and locomotion problems, strong activation of glial cells, lack of trophic support, mitochondrial-related alterations, and other molecular features in the cerebellum of YG8-800 mice that closely resemble what occurs in humans. These findings provide valuable information that could be used as clinical readouts in future studies in which new therapeutic approaches or targets are tested.

## Conclusions

As summarized in Fig.8, by establishing and characterizing this new humanized FRDA murine model, we have demonstrated that FXN-deficient YG8-800 mice show a progressive reduction of the main cerebellar neuronal population, the CGNs, which correlates with progressive impairment in the locomotor function of these mice. Moreover, our studies revealed an early activation of astrocytes and microglia, which are possibly contributing to the observed neuroinflammation and neurodegeneration. This in-depth characterization of the YG8-800 mouse model allows the investigation of the pathogenic molecular mechanisms responsible for FRDA, representing as well a convenient experimental tool for the development of new therapies that could limit the progression of the disease.

**Figure 8:**
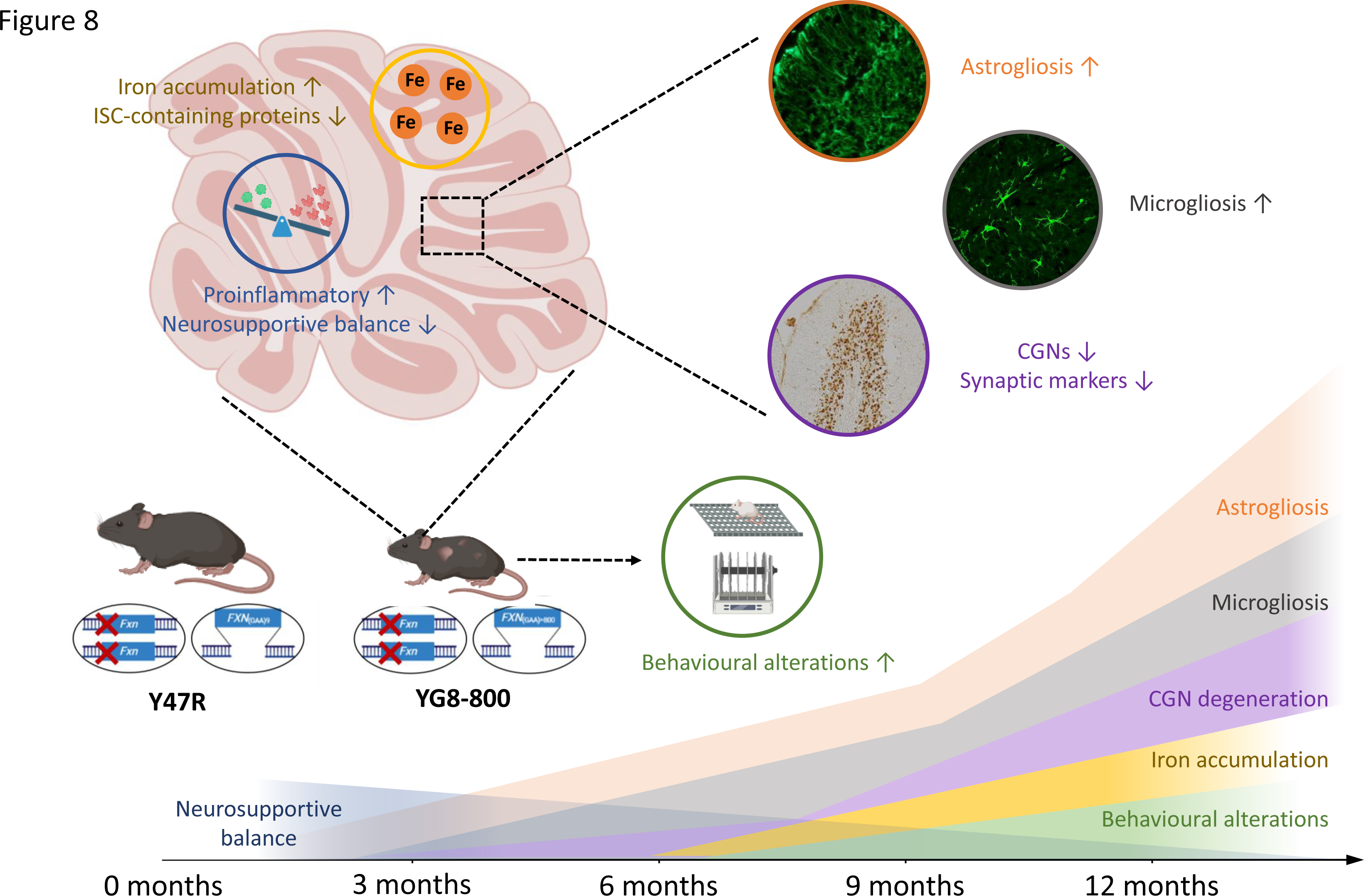
Diagram showing the main findings identified in the YG8-800 murine model of FRDA. Schematic representation showing the chronology of altered locomotor behavior and biochemical and histological changes observed in the cerebellum of FXN-deficient mice over 12 months.

## List of abbreviations

ARCAs: Autosomal recessive cerebellar ataxias
CGN: Cerebellar granule neurons
CNS: Central nervous system
ETC: Electron transport chain
FRDA: Friedreich’s ataxia
FXN: Frataxin
GAA: Guanine-adenine-adenine
GL: Granular layer
GM: Gray matter
ISC: Iron-sulfur clusters
ML: Molecular layer
PC: Purkinje cell
WM: White matter
YAC: Yeast artificial chromosome

## Declarations

### Ethics approval and consent to participate

Not applicable

### Consent for publication

Not applicable.

### Availability of data and material

All data used and analyzed for the current study are available from the corresponding author upon reasonable request.

### Contributions

Data curation: AVA, JGC, SHM; Formal analysis: AVA, JGC, SHM, MRP, FL; Funding acquisition: JDN, FL, SHM. Investigation: AVA, JGC, SHM, MRP, MAL; Methodology: AVA, JGC, SHM, MRP, FL; Project administration and supervision: FL; Resources: JDN, FL; Visualization: AVA; Writing –original draft preparation: AVA., SHM, JGC, FL; Writing –review & editing: AVA., SHM, JGC, MRP, JDN, FL.

## Acknowledgements

The authors are grateful to the Confocal Microscopy, Flow Cytometry, and Genomics scientific facilities of the CBM Severo Ochoa for their excellent assistance. We thank Elia Pérez for the valuable discussion on statistics.

---

Additional file 1: Table S1. List of qPCR primers.

Additional file 2: Movie S1. Y47R and YG8-800 mice were recorded at 9 months of age. The video shows the smaller size and motor impairments in YG8-800 mice compared to Y47R.

