## Additional Table S1 for "Glial cell activation precedes neurodegeneration in the cerebellar cortex of the YG8-800 murine model of Friedreich’s ataxia"

**Table S1**. Human/mouse primer sequences used for qPCR

| **Protein (Gene name)B2:D38B47B2:C32B2:D40BB2:D40** | **Primer Name** | **Sequence 5' -> 3'** |
|---|---|---|
| Arginase 1 (*Arg1*) | m-Arg1 (Fw) | CAGAAGAATGGAAGAGTCAG |
| | m-Arg1 (Rv) | CAGATATGCAGGGAGTCACC |
| Brain derived neurotrophic factor (*Bdn* f) | m-Bdnf (Fw) | GGCTGACACTTTTGAGCACGTC |
| | m-Bdnf (Rv) | CTCCAAAGGCACTTGACTGCTG |
| Fibroblast growth factor 2 (*Fgf2*) | m-Fgf2 (Fw) | CACCAGGCCACTTCAAGGA |
| | m-Fgf2 (Rv) | GATGGATGCGCAGGAAGAA |
| Frataxin (*FXN*) | h-FXN (Fw) | TGGAATGTCAAAAAGCAGAGT |
| | h-FXN (Rv) | CCACTCCCAAAGGAGACATC |
| Frataxin (*FXN*) for genotyping | h-FXNgen (Fw) | CCCCTGATTTGCTGTATGCT |
| | h-FXNgen (Rv) | CTCAAGGTCTCCGCACTTG |
| Glyceraldehyde-3-phosphate dehydrogenase (*Gapdh*) | m-Gapdh (Fw) | CCCCTGGCCAAGGTCATCCATG |
| | m-Gapdh (Rv) | CAGTGAGCTTCCCGTTCAGCTC |
| Glial cell-derived neurotrophic factor (*Gdnf*) | m-Gdnf (Fw) | AAAGTAGGCCAGGCATGTTG |
| | m-Gdnf (Rv) | TTCGCACTGTAGCAGGAATG |
| Solute carrier family 1 member 3 (*Glast/Slc1a3*) | m-Glast (Fw) | GCGATTGGTCGCGGTGATAATG |
| | m-Glast (Rv) | CGACAATGACTGTCACGGTGTAC |
| Solute carrier family 1 member 2 (*Glt1/Slc1a2*) | m-Glt1 (Fw) | TGGACTGGCTGCTGGATAGA |
| | m-Glt1 (Rv) | CGGTGTTGGGAGTCAATGGT |
| Glutamate-ammonia ligase (*Glul*) | m-Glul (Fw) | CTGCCATACCAACTTCAGCACC |
| | m-Glul (Rv) | CTGGTGCCTCTTGCTCAGTTTG |
| Hepatocyte growth factor (*Hgf*) | m-Hgf (Fw) | CATTGGTAAAGGAGGCAGCTATAAA |
| | m-Hgf (Rv) | GGATTTCGACAGTAGTTTTCCTGTAGG |
| Insulin-like growth factor binging protein 2 (*Igfbp2*) | m-Igfbp2 (Fw) | CAGACGCTACGCTGCTATCC |
| | m-Igfbp2 (Rv) | CTCCCTCAGAGTGGTCGTCA |
| Insulin-like growth factor binging protein 3 (*Igfbp3*) | m-Igfbp3 (Fw) | CCTCAATGTGCTGAGTCCCAGA |
| | m-Igfbp3 (Rv) | CTTGTCCACACACCAGCAGAAG |
| Interleukin 1 alpha (*Il1a*) | m-Il1a (Fw) | GCACCTTACACCTACCAGAGT |
| | m-Il1a (Rv) | AAACTTCTGCCTGACGAGCTT |
| Interleukin 1 beta (*Il1b*) | m-Il1b (Fw) | TGCCACCTTTTGACAGTGATG |
| | m-Il1b (Rv) | TGATGTGCTGCTGCGAGATT |
| Interleukin 6 (*Il6*) | m-Il6 (Fw) | TACCACTTCACAAGTCGGAGGC |
| | m-Il6 (Rv) | CTGCAAGTGCATCATCGTTGTTC |
| Nitric oxide synthase 2 (*iNos/Nos2*) | m-iNos (Fw) | TGCATGGACCAGTATAAGGCAAGC |
| | m-iNos (Rv) | GCTTCTGGTCGATGTCATGAGCAA |
| Potassium inwardly rectifying channel subfamily J member 10 (*Kir4.1/Kcnj10*) | m-Kir4.1 (Fw) | TGCGGAAGAGTCTCCTCATTGG |
| | m-Kir4.1 (Rv) | GTCTGAGGCTGTGTCTACTTGG |
| Nerve growth factor (*Ngf*) | m-Ngf (Fw) | CATGGGGGAGTTCTCAGTGT |
| | m-Ngf (Rv) | GCACCCACTCTCAACAGGAT |
| Ribosomal RNA 18S (*Rna18S*) | m-Rna18S (Fw) | GCAATTATTCCCCATGAACG |
| | m-Rna18S (Rv) | GGGACTTAATCAACGCAAGC |
| Transforming growth factor beta 1 (*Tgfb1*) | m-Tgfb1 (Fw) | TGATACGCCTGAGTGGCTGTCT |
| | m-Tgfb1 (Rv) | CACAAGAGCAGTGAGCGCTGAA |
| Tumor necrosis factor (*Tnfa*) | m-Tnfa (Fw) | CCCTCACACTCAGATCATCTTCT |
| | m-Tnfa (Rv) | GCTACGACGTGGGCTACAG |
| Neurotrophic receptor tyrosine kinase 2 (*Trkb/Ntrk2*) | m-Trkb (Fw) | GAATAACGGAGACTACACCCTGATG |
| | m-Trkb (Rv) | ACCACAGATGCAATCACCACC |
